# Warm temperature modifies cell fates to reduce stomata production in Arabidopsis

**DOI:** 10.1101/2024.12.19.629334

**Authors:** Josué Saiz-Pérez, Alexandra Baekelandt, Jonatan Illescas-Miranda, Lieven Sterck, Marnik Vuylsteke, Eun-Ji Kim, Boyu Guo, Bénédicte Desvoyes, Crisanto Gutierrez, Eugenia Russinova, Carmen Fenoll, Montaña Mena

**Affiliations:** Facultad de Ciencias Ambientales y Bioquímica, Universidad de Castilla-La Mancha, Av Carlos III sn, 45071, Toledo, Spain; Department of Plant Biotechnology and Bioinformatics, Ghent University, Technolgiepark 71, Ghent 9052, Belgium; Center for Plant Systems Biology, VIB, Technolgiepark 71, Ghent 9052, Belgium; Centro de Biología Molecular Severo Ochoa, CSIC-UAM, Nicolás Cabrera 1, Cantoblanco, 28049, Madrid, Spain

**Keywords:** warm temperature, stomatal lineage development, SPCH, MUTE, PIF4, transcriptome, cell fate

## Abstract

- Stomatal abundance decrease in Arabidopsis triggered by warm-temperature is attributed to PIF4-mediated repression of *SPEECHLESS* (*SPCH)* expression. We identified the unknown developmental and transcriptional basis of this adaptive response.
- We traced stomatal lineages in vivo using cell-identity marker lines and mutants, quantified epidermal traits and conducted RNA sequencing under oscillating temperatures.
- Prolonged warm-temperature or PIF4-overexpression altered cell-fates, inducing diverted stomatal precursors (DPs) that lacked stomatal fate, accounting for stomata reduction. DPs originated from meristemoids that lost *SPCH* expression, lacked *MUTE* expression and exited the cell cycle. Short warm-temperature pulses allowed later recovery of *SPCH* expression, and did not induce DPs or stomata reduction. Comparison of transcriptomes obtained during warm-temperature pulses with stomatal lineage cell-specific profiles identified gene expression changes and contrasted their reversibility. Though warm-temperature silenced key stomatal drivers, most lineages formed stomata through partly modified transcriptional landscapes that promoted uncommitted cell identities and included alternative pathways.
- Expression changes in stomatal regulators and cell-fate changes explain lineage progression under fluctuating temperatures. Since short-term temperature oscillations prevail in natural conditions, the requirement of long warm-temperature exposure to trigger DPs would prevent stomata reduction by occasional temperature rises. Promoting uncommitted lineage stages provides flexibility to stomatal development under environmental changes.

## INTRODUCTION

Gas exchange between the plant and the atmosphere takes mostly place through stomata (Ziegler, 1987). These valves mediate CO_2_ uptake and the H_2_O vapor loss that drives root water and nutrient uptake and refrigerates the plant, critically influencing growth and reproduction (Raven, 2002). Stomata abundance determines the maximum area available for gas exchange (Franks *et al*., 2009; Dow *et al*., 2014), its adaptive value exemplified by broad inter- and intraspecific natural variation, and by its adjustment under different environmental conditions (Woodward, 1987; Woodward *et al*., 2002; Delgado *et al*., 2011; Dittberner *et al*., 2018; Bertolino *et al*., 2019).

In *Arabidopsis thaliana*, stomata form from protodermal cells that initiate stomatal cell lineages, whose cell divisions and differentiation events are determined by endogenous programs driven by the transcription factors SPEECHLESS (SPCH), MUTE and FAMA, and modulated by environmental cues (Bergmann & Sack, 2007; Pillitteri & Torii, 2012). A stomatal lineage is initiated when SPCH accumulates in a meristemoid mother cell, driving an asymmetric cell division (ACD) that generates a smaller, self-renewing meristemoid, (MacAlister *et al*., 2007), and a larger stomatal lineage ground cell (SLGC) (Pillitteri & Torii, 2012). SLGCs differentiate as pavement cells, constituting most of the epidermal surface (Geisler *et al*., 2000). After several ACDs, the late-meristemoid acquires guard mother cell (GMC) identity triggered by MUTE, which orchestrates a symmetric cell division (SCD) to produce the guard cell (GC) pair (Pillitteri *et al*., 2007; Han *et al*., 2018). FAMA drives GC differentiation (Ohashi-Ito & Bergmann, 2006; Hachez *et al*., 2011) and ensures a single SCD (Lai *et al*., 2005; Xie *et al*., 2010; Lee *et al*., 2014). SPCH, MUTE and FAMA, interacting with SCRM/2 partners, influence stomata abundance (Kanaoka *et al*., 2008). As stomatal movements need surrounding non-stomatal cells, robust mechanisms prevent that stomata are formed in contact, based on inhibition of SPCH and MUTE activities in cells contacting meristemoids/stomata. Meristemoids secrete inhibitory peptides, perceived by membrane receptors containing ERECTA family receptor-kinases that activate MAPK cascades led by YODA, whose outcome is marking SPCH, SCRM/2 and/or MUTE for degradation (Lee & Bergmann, 2019; Herrmann & Torii, 2021). These repressive signaling pathways ensure functional patterns and, interacting with hormonal and environmental signals, modulate stomata abundance to optimize plant performance (Han & Torii, 2016; Qi & Torii, 2018).

Among the cues shaping stomatal development are warm temperatures (warm-T). In Arabidopsis, temperatures slightly above the optimum induce thermomorphogenesis (Erwin *et al*., 1989), reprogramming developmental networks and modifying plant architecture (Balasubramanian *et al*., 2006; Crawford *et al*., 2012; Quint *et al*., 2016; Casal & Balasubramanian, 2019; Vu *et al*., 2019). Thermomorphogenesis includes adaptative mechanisms to enhance plant fitness under high temperatures, many involving PHYTOCHROME INTERACTING FACTOR 4 (PIF4) (Gray *et al*., 1998; Koini *et al*., 2009; Franklin *et al*., 2011; Quint *et al*., 2016; Delker *et al*., 2022). In Arabidopsis (Col-0), warm-T triggers a moderate reduction in stomatal abundance (Crawford *et al*., 2012; Pérez-Bueno *et al*., 2022) linked to a transcriptional negative feedback loop between PIF4 and SPCH (Lau *et al*., 2018). However, virtually nothing is known about how cell divisions and identities in stomatal lineages are altered during the exposure to warm-T that activates this SPCH-PIF4 repressive loop, and which are the subjacent molecular changes.

In this work, we analyzed stomatal lineage progression at warm- and control-T combining live-cell imaging and lineage-cell tracing with epidermal phenotyping, genetics and transcriptomics. Warm-T or PIF4 overexpression caused a fraction of stomatal precursors to lose their identity and become diverted, explaining stomatal index reduction, and hinting at developmental mechanisms underlying lineage progression. Triggering diverted precursor fate required extended exposure to warm-T, below which the process is reversible and stomatal index remains unchanged. Despite heat-induced gene reprogramming silenced key positive drivers of stomatal development as SPCH and MUTE, most lineages progressed and formed stomata. Transcriptomics revealed that warm-T shifted lineages towards uncommitted cell stages, which regained committed fates during recovery at control temperature. This indicates that stomatal development under changing temperatures occurs through partly rewired gene circuits involving alternative pathways.

## MATERIALS AND METHODS

### Plant materials and growth conditions

*Arabidopsis thaliana* (L.) Col-0 accession (N1092) was obtained from Nottingham Arabidopsis Stocks Centre. *pif4-2* and *pifQ* (*pif1.3.4.5*) were donated by Salome Prat (Bernardo-García *et al*., 2014; Murcia *et al*., 2022). The *iMUTEmute* line, carrying a *MUTE* expression system for conditional complementation of the loss-of-function *mute-3* mutation, was described by Triviño *et al*. (2013). Conditional overexpressors *iPIF4oe* and *iSPCHoe* under the constitutive pG10-90 promoter (Ishige *et al*., 1999) were derived from pER8 (Zuo *et al*., 2000) in the TRANSPLANTA consortium (Coego *et al*., 2014). The plasma membrane marker line *AtML1pro:mCherry-RCl2A* was published by Kim *et al*. (2023). *TMMpro:GFP-GUS* and *flp-1* was a gift from Fred Sack (Nadeau & Sack, 2002). *FLPpro:GUS-GFP* was published by Wang *et al*. (2015). To obtain *MUTEpro:GFP*, a 2 kb-long upstream sequence from the *MUTE* start codon was cloned into the Gateway compatible entry vector pDONRp4-p1r (primers listed in Supplemental Table S1). GFP cDNA was subcloned into pDONR221 and both recombined in pK7m24GW,3 destination vector. Plant transformation was performed by *Agrobacterium tumefaciens-*mediated floral dip (Clough & Bent, 1998), and transgenic seedlings selected in MS plates with 50 mg/l kanamycin. Dual reporters for *SPCHpro:GFP* (de Marcos *et al*., 2017)*, MUTEpro:GFP* and *TMMpro:GFP-GUS* carrying *AtML1pro:mCherry-RCl2A* were generated through sexual crosses. The *PlaCCI Lti6b-GFP* line carrying cell cycle and plasma membrane markers was a gift from Keiko Torii (Desvoyes *et al*., 2020; Zuch *et al*., 2023).

Seeds were sterilized with chlorine gas (Clough & Bent, 1998) and stratified for 4 days at 4°C prior to light exposure. Seedlings were grown in ½ strength MS and 1% (w/v) agar. Conditional expression was induced with 10 µM of β-estradiol in DMSO. All experiments were in vitro unless otherwise indicated. Growth chambers were set at 70% RH, 70 µmol·m^−2^·s^−1^ and long day (16-h-light/8-h-dark). Control temperature was 21°C and warm temperature 28°C. For soil-grown plants, 7mm peat pellets (Jiffys©) were used with the same conditions. To avoid a germination-selection bias, seedlings were grown for 3 days at 21°C prior to 28°C exposure.

### Quantitative stomatal parameters

Cell proportions and sizes were determined in abaxial cotyledon or leaf epidermis from micrographs with Fiji/ImageJ (Schneider *et al*., 2012). Cell types were estimated by morphological criteria, marker expression or cell fate. Stomatal index (SI) is the number of stomata/number of total cells × 100. Diverted precursor index (DPI) is the number of diverted precursors/number of total cells × 100. Stomatal density is the number of stomata/mm^2^. In mature organs, cells were counted in 0,4mm^2^ of the mid-central abaxial epidermis. In developing organs, lineage index (LI) is [number of stomata + number of meristemoids]/number of total cells × 100, meristemoid index is the number of meristemoids/number of total cells × 100. An area of 0,18 mm^2^ was examined for developing organs. Percentage of fluorescent marker expression is number of expressing cells/total number of cells × 100. Statistical analyses were performed using SPSS (IBM Inc). Figure legends detail numbers of plants or cells scored in different experiments and statistical tests.

### Microscopy

For differential interference contrast microscopy (DIC) imaging, organs were hand-excised and fixed in ethanol:acetic acid 9:1 (v/v) for 16 hours, rehydrated by serial 1-hour incubation at room temperature in ethanol of 90%, 70%, 50%, 30% (v/v) dilutions followed by distilled water, and cleared in chloral hydrate:glycerol:water (8:2:1 w/v/v). Micrography was performed under a Nikon Eclipse 90i microscope, using a Gryphax (JenoptikTM) camera. For leaf lineage tracing, nail polish replicas of serial dental resin imprints (Kagan *et al*., 1992) were analyzed with a Hitachi TM-1000 scanning electron microscope.

Confocal images were taken with a Leica SP8 spectral microscope and processed with Fiji/ImageJ. Cell outlines were viewed with the plasma membrane marker *AtML1pro:mCherry-RCl2A* or by staining with 50 µg·ml^−1^ propidium iodide. For timelapse imaging and lineage tracing, seedlings were transferred to sterilized ½ strength MS, 1% agar media mounted in a Lab-Tek™ II chamber (Thermo Scientific) kept at 21°C. Z-stacks through the epidermis were captured every 30 minutes for timelapse imaging. For sequential time-course tracing, after each recording seedlings were transferred back to media plates and kept under the established temperature.

### Ploidy level assessment

21 days-old *AtML1pro:mCherry-RCl2A* cotyledons were stained with 0.5 mg ml^−1^ 4,6-diamidino-2-phenylindole dihydrochloride (DAPI) for 5 minutes, rinsed with distilled water and the abaxial epidermis examined by confocal microscopy. Nuclei projected area was obtained and measured using Fiji/ImageJ. Ploidy levels associated to nuclei size were estimated following a size-ploidy range as described (Triviño *et al*., 2013).

### RNA extraction and transcriptome analysis

Total RNA was extracted from 3–4-day-old seedlings after temperature treatments using Promega Total RNA isolation kit. The three biological replicates per treatment contained 50 seedlings and temperature shifts considered the Zeitgeber time. Library construction and sequencing was carried out by Macrogen Inc. through TruSeq Stranded mRNA Library Prep Kit (Ilumina, Inc.) and paired-end sequencing. Quality of 150bp length reads was assessed with FastQC (Wingett & Andrews, 2018). Low quality reads and adaptor sequences were filtered out using Trimmomatic (Bolger *et al*., 2014). Raw reads were aligned to reference transcriptome ARAPORT11 and quantified using Salmon (Patro *et al*., 2017). Transcripts were summarized using Tximport (Soneson *et al*., 2016) and differential gene expression calculated with EdgeR (McCarthy *et al*., 2012). Differentially expressed (DE) genes were statistically tested by false discovery rate (FDR) algorithm. Reference data are deposited under GEO accession no. GSE278519.

### RT-qPCR analysis

RNA samples obtained for RNAseq were used for cDNA synthesis with qScript cDNA SuperMix (Quantabio) following manufactureŕs instructions. SYBR green qPCR master mix (Thermo Scientific) was used for real-time-quantitative PCR (RT-qPCR) analysis in a LightCycler 480 II instrument (Roche Diagnostics), with primers listed in Supplemental Table S1. *ACT2* (AT3G18750) and *UBQ10* (AT4G05320) were used as references. Relative expression and normalization were calculated over sampling timepoints using the Taylor *et al*. (2019) reference method.

## RESULTS

### Warm temperature induces diverted stomatal precursors that correlate with stomata reduction

In Arabidopsis (Col-0), growth at warm-T (28°C) decreases the stomatal index in mature cotyledons (Lau *et al*., 2018). To explore the developmental basis of this decrease, we examined cell morphology in the abaxial cotyledon epidermis of plants grown at 28°C for 19 days (Fig. **1a**). Along stomata and pavement cells, we found groups of cells with the relative disposition of stomatal lineages that, however, remained under-expanded and lacked a central stoma. Instead, in the spiral arrangements the cell occupying the expected stoma location had uncommon shape and larger size than meristemoids (colored in orange in Fig. **1a**). We presumed that these atypical central cells were stomatal precursors that lost their fate, and termed them “diverted stomatal precursors” (DPs).

**Fig. 1.**
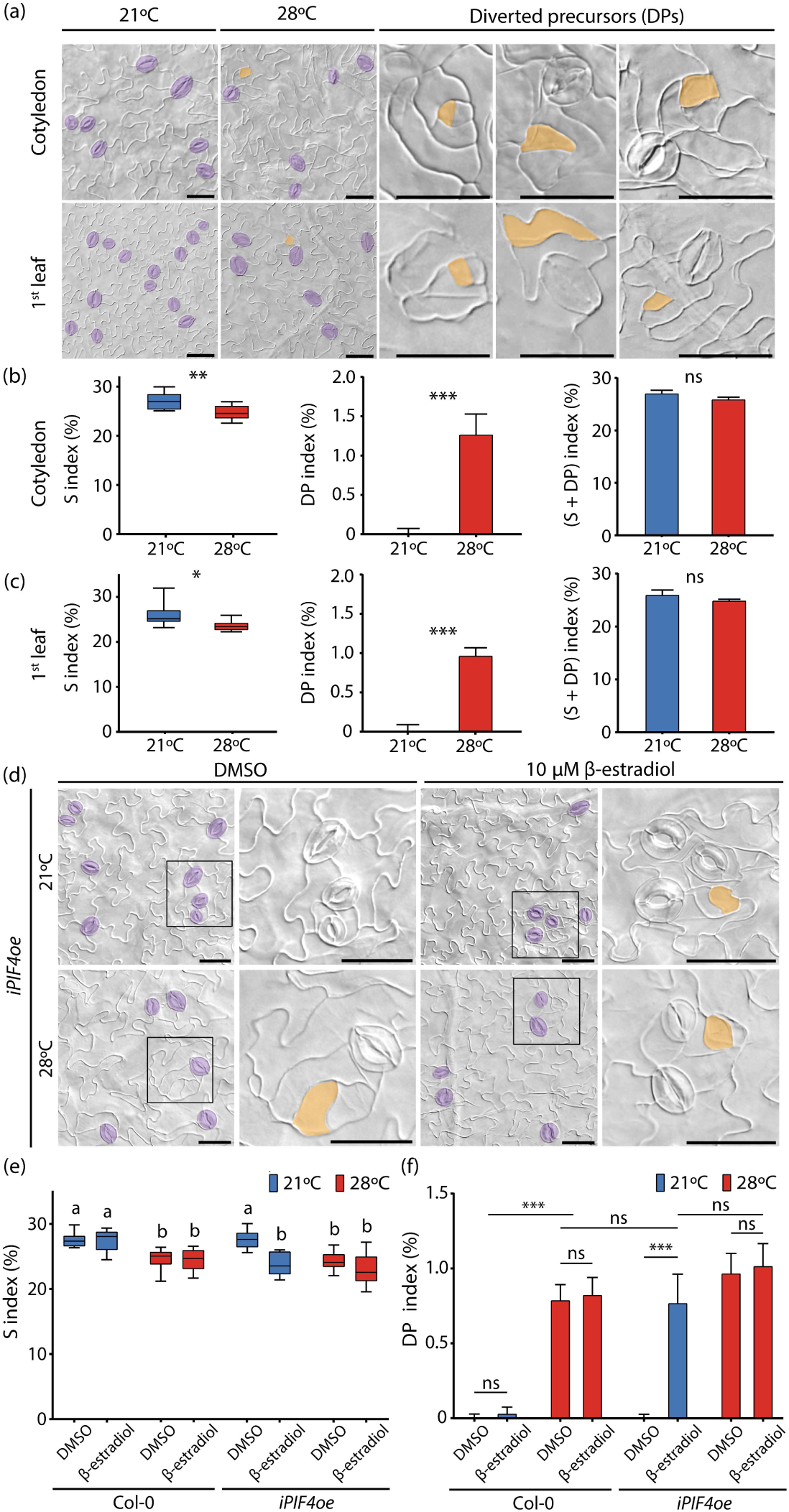
Warm temperature or PIF4 overexpression induce diverted stomatal precursors and reduce stomatal index. (a,d) Representative abaxial DIC images of fully developed cotyledons and 1^st^ leaves of 21-days-old Col-0 (a) grown at control (21°C) and warm (28°C) temperatures, and *iPIF4oe* (d) supplemented with DMSO (control) or 10 µM β-estradiol (induced). Stomata are colored in purple and diverted precursors in orange. Scale bar = 50 µm. (b,c,e,f) Abaxial stomatal (S), diverted precursor (DP) and (S+DP) index are shown for Col-0 cotyledons (b) and 1^st^ leaves (c) in plants grown at 21°C and 28°C. (e,f) Cell indexes in *PIF4oe* cotyledons from non-induced (DMSO) and induced (10 µM β-estradiol treated) plants. (b, c, e) Box-plots show median, interquartile range 1-3 and the maximum-minimum values. (b, c, f) Histograms show the mean ± SEM. (b,c) Asterisks represent *P* < 0.05 (*), *P* < 0.01 (**), *P* < 0.001 (***) and ns indicates not significant differences according to a Student’s t-test (b, c) or a Kruskal-Wallis pairwise comparison test (f). (e) Letters indicate statistical differences according to one-way ANOVA with post hoc Tukey HSD. (b, c, e, f) *n* = 10 plants per genotype and treatment.

In cotyledons, DPs represented 1.5% of total epidermal cells as measured by the “diverted precursor index” (DPI; Fig. **1b**), and potentially depleted 4.5% of the initiated lineages for stomatal production. Despite a low occurrence, DP numbers consistently account for the SI reduction observed at warm-T. Considering that each DP could have yielded a stoma, we calculated the (S+DP) Index, which did not differ from the SI at 21°C as control-T (Fig. **1b**). DPs were not a singularity of the embryonic organ, as they appeared in the first leaf in a similar proportion (Fig. **1a,c**), suggesting that warm-T reduced stomatal numbers by making a fraction of stomatal precursors withdraw from stomatal fate.

### PIF4 is necessary and sufficient for DP formation

Since the stomatal response to warm-T was ascribed to PIF4 (Lau *et al*., 2018), we examined mature abaxial cotyledons of *pif4-2* and *pif 1-3-4-5* (*pifQ*) plants grown at 21°C or 28°C (Fig. **S1**). The mutants showed no SI reduction by warm-T, which triggered a 3.5% SI reduction in Col-0. DPs were never observed in *pif4-2* nor *pifQ*. The *pif4-2* phenotype indicated that DP induction by warm-T is PIF4-dependent and confirmed that PIF4 is required for the temperature-dependent SI decrease. *pifQ* demonstrated that PIF4 has non-redundant functions with PIF1/3/5 on the thermal control of stomatal development.

If *PIF4* transcriptional activation in meristemoids by warm-T underlie DP formation, *PIF4* over-expression should result in DP production. Using a β-estradiol-inducible *PIF4* over-expressor line (*iPIF4oe*) we determined SI and DPI in 21-days-old cotyledons grown with or without β-estradiol at both temperatures (Fig. **1d-f**). At 21°C, β-estradiol-induced *iPIF4oe* cotyledons showed the same DPI and SI than Col-0 grown at 28°C, indicating that increased PIF4 levels and warm-T produced similar fate defects on stomatal precursors. PIF4 overexpression did not enhance the SI reduction caused by warm-T, as induced *iPIF4oe* cotyledons had the same SI and DPI at 21°C and 28°C (Fig. **1e,f**). Thus, DPs can form at control-T by overproducing PIF4, supporting that their appearance underlies the PIF4-dependent SI reduction observed at warm-T. The response seems saturated by the PIF4 levels induced by warm-T in meristemoids described by Lau et al, (2018), as ubiquitous PIF4 overexpression had no additional effect.

### Diverted cells are miss-fated stomatal precursors

We hypothesized that the atypical cells appearing at warm-T or upon PIF4-overexpression were stomatal precursors with diverted fate. To assess this, we traced meristemoid histories using a *SPCHpro:GFP* line, monitoring meristemoid fate (stoma, meristemoid or DP) and *SPCHpro* expression every 48h. This was done after moving 3-day-old seedlings from 21°C to 28°C or maintaining them at 21°C (Fig. **2a**). Consistent with previous reports (Lau *et al*., 2018), growth for 48h at warm-T greatly reduced proportions of *SPCHpro*-expressing meristemoids (Fig. **2b**), while at 21°C most expressed *SPCHpro*. By 96h, lineages had progressed and most meristemoids lost *SPCHpro* expression. A fraction of the meristemoids that after 48h at 28°C showed no *SPCHpro* signal, by 96h displayed DP morphology (Fig. **2a**). These data confirm that warm-T-induced DPs derive from meristemoids that lost *SPCH* expression, and that *SPCH* silencing seems necessary for their loss of stomatal fate.

**Fig. 2.**
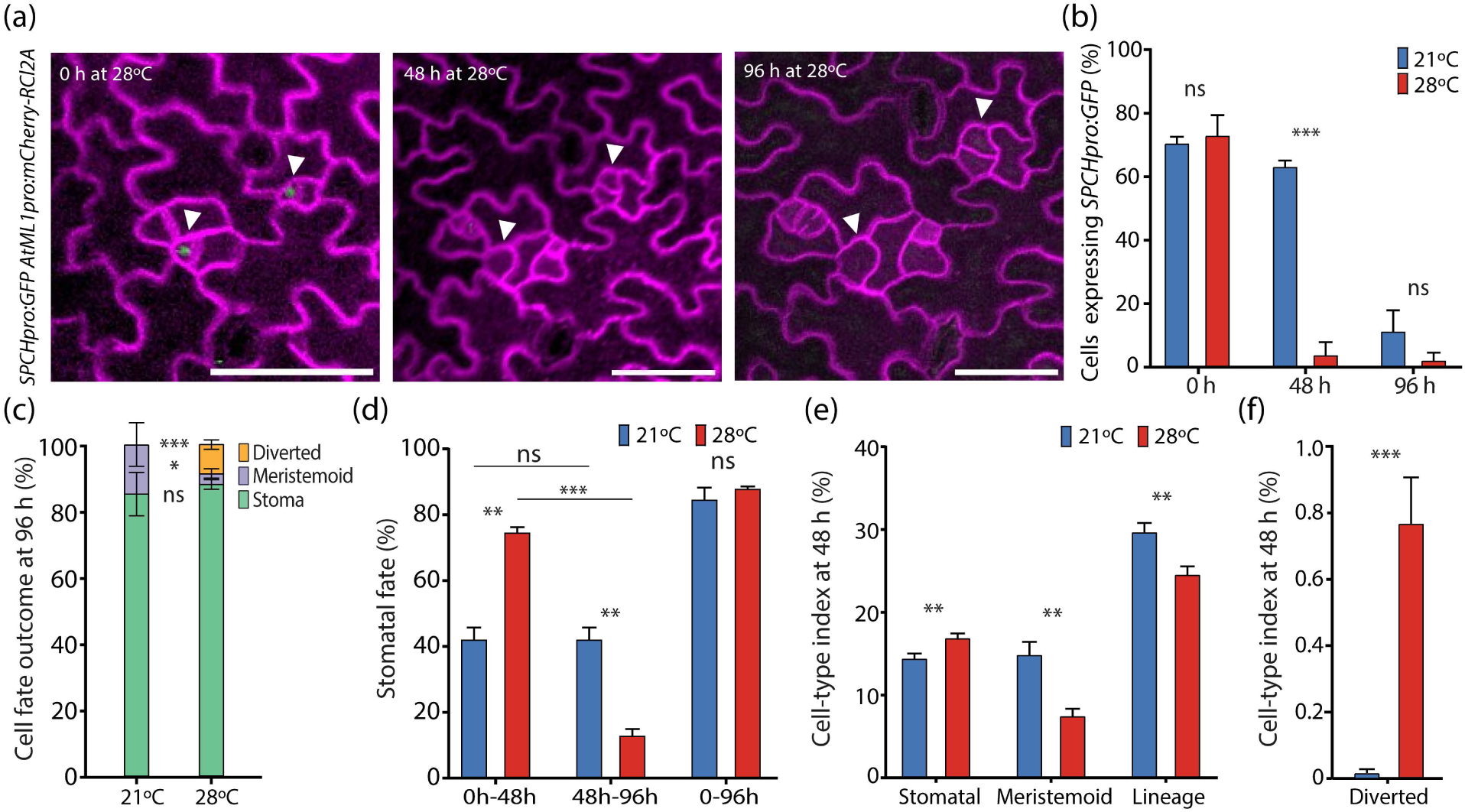
Diverted precursors derive from meristemoids. (a) Representative confocal images of in vivo *SPCHpro:GFP* transcriptional activity dynamics in traced stomatal lineages. Tracing started in the abaxial epidermis of 3 days-old-cotyledons (0h) and individual lineages were recorded after 48h and 96h of exposure to warm-T. White arrowheads indicate meristemoids with *SPCHpro:GFP* signal (green) that resulted in diverted precursors. Cell contours (purple) show the plasma membrane marker *AtML1pro:mCherry-RCl2A.* Scale bar = 50 µm. (b) Percentage of traced cells expressing *SPCHpro:GFP* at 0h, 48h and 96h after transfer to 21°C or 28°C. (c) Percentage of meristemoids, stomata and diverted precursors at 96h. (d) Stomatal commitment timing scored in traced lineages at intervals 0h-48h, 48h-96h, and 0h-96h at 21°C or 28°C. Stomatal, meristemoid and total lineage indexes (e), and diverted precursor index (f) at 48h in abaxial cotyledons grown at 21°C and 28°C. Error bars represent SEM. Asterisks represent *P* < 0.05 (*), *P* < 0.01 (**), *P* < 0.001 (***) and not significant (ns) differences (Student’s t test) between temperature treatments. Individual stomatal lineages (*n* = 120) from independent plants (*n* ≥ 3) and treatments were followed by time-course confocal imaging.

Individual meristemoid tracing revealed that approximately 85% had formed stomata by 96h at either temperature. But at 28°C the remaining meristemoid number was lower than that at control-T, and 8% had produced DPs (Fig. **2c**). After 48h at 28°C almost all meristemoids failed to express *SPCHpro*, but by 96h most had formed stomata. This implies that, surprisingly, *SPCHpro* silencing is not sufficient to drive meristemoids into non-stomatal fate, perhaps because of residual SPCH activity (see below). At warm-T almost 80% of the meristemoids had formed stomata by 48h (Fig. **2d**) compared to barely 40% at control-T, indicating that warm-T accelerates stomatal development in most lineages, while triggering diverted fates in only a small fraction. At 21°C, differentiation was progressive over the 96h supervised, with 40% of meristemoids producing a stoma every 2 days, whereas at 28°C these were 75% during the first 48h. By that time, differences in cell-transition timing resulted in lower meristemoid index and higher SI at 28°C than at 21°C (Fig. **2e**). As some meristemoids were diverted at 28°C (Fig. **2f**), lineage index (LI) was lower at 28°C than at 21°C (Fig. **2e**), indicating that 48h at warm-T decreased the meristemoid population potential to form stomata.

*PIF4*-overexpression at control-T produced DPs and reduced SI very early during cotyledon development (Fig. **S2**). Induced 5-days-old *iPIF4oe* showed both SI and LI decreases compared to non-induced plants, and accumulated abundant DPs. These broader and faster responses than warm-T-treated Col-0 plants of the same age (3 days plus 48h; Fig**. 2e**) could be explained because Col-0 seedlings are kept at 21°C for the first 3 days before exposure to 28°C, whereas *iPIF4oe* was induced since germination. Therefore, lineage cells in *iPIF4oe* plants were exposed to high PIF4 levels from earlier developmental stages and for a longer time. These results further strengthen the interpretation that PIF4 mediates DP production at warm-T (Fig. **1**).

Warm-T also downregulated *SPCHpro:GFP* in leaf primordia (Fig. **S3 a-c**) and we traced DPs back to meristemoid-shaped cells through serial imprints (Fig. **S3 d-e**). Thus, warm-T induces a small but significant and consistent proportion of meristemoids in the young organs to become diverted.

### Alteration of SPCH levels interferes with DP production and SI reduction

The finding that a stabilized SPCH repressed *PIF4* transcription lead to propose a reciprocal negative feedback loop that maintains enough SPCH for lineage completion at warm-T (Lau *et al*., 2018), but the impact of SPCH and PIF4 levels on the SI reduction triggered by warm-T was not assessed. We tested if elevating *SPCH* expression influenced SI and DP production using a β-estradiol-dependent *SPCH* overexpressing line under a constitutive promoter, *iSPCHoe* (Fig. **S4**). Seedlings were grown under induced and non-induced conditions at control-T or warm-T, and SI and DPI assessed in 21-days-old cotyledons. Uninduced plants showed the expected SI reduction at 28°C and produced DPs. Induced plants did not reduce their SI at warm-T and their DP production was residual, implying that, as predicted (Lau *et al*., 2018) higher SPCH levels counteract PIF4 effects. The residual DPs in induced plants was statistically similar at control and warm-T, indicating that constitutive high SPCH levels had a minor but detectable effect on stomatal precursors fate.

### *MUTE* expression prevents diverted precursors and is silenced by warm temperature

*MUTE* expression in meristemoids mark their transition to GMCs, committed to form stomata. To study how DP fate relates to *MUTE* expression, we used a *MUTEpro:GFP* to trace meristemoids transitioning to GMCs in cotyledons exposed to 28°C during 48h and 96h (Fig. **3a**), finding that warm-T silenced *MUTEpro* almost completely. By tracking back lineage history of stomata and DPs identified at 96h, we found that none of the meristemoids that became DPs had previously expressed *MUTEpro:GFP.* Conversely, all precursors that expressed *MUTEpro:GFP* formed stomata (Fig. **3d,e**), implying that once GMC identity is established by MUTE, commitment to stomatal fate becomes insensitive to warm-T, and that stomatal precursors diversion takes place prior to the onset of *MUTE* expression. Puzzlingly, and as was the case with *SPCHpro*, regardless the profound *MUTEpro* silencing at warm-T (Fig. **3b,c**), most lineages were able to form stomata (see below).

**Fig. 3.**
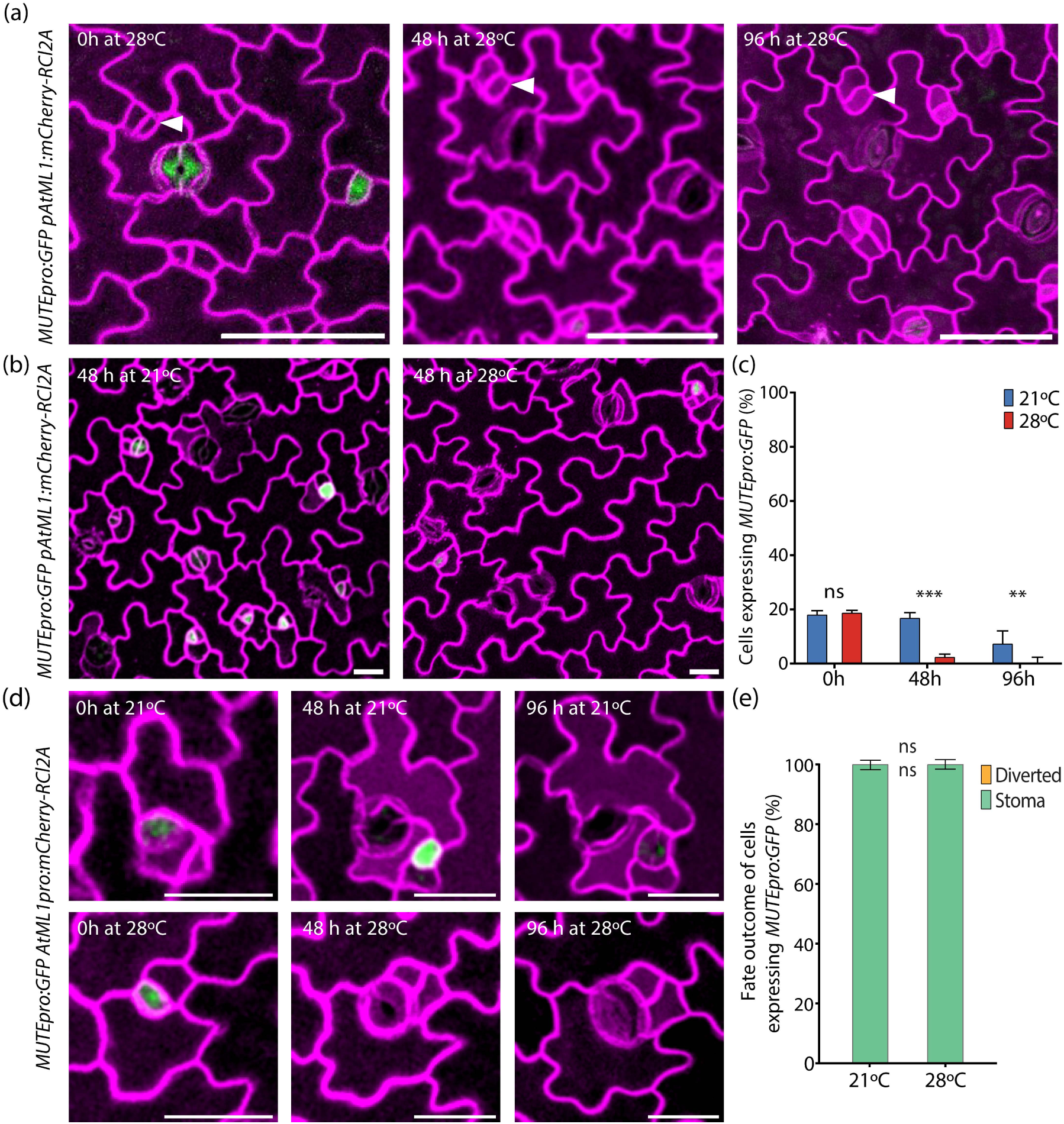
*MUTE* is silenced by warm temperature and prevents diverted precursor fate. (a) Confocal in vivo tracing of *MUTEpro:GFP* expression in 3 days-old abaxial cotyledons (0 hours) exposed to 28°C during 48 and 96 hours. White arrowhead identifies diverted precursors. Cell contours (purple) show the plasma membrane marker *AtML1pro:mCherry-RCl2A.* (b) Representative *MUTEpro:GFP; AtML1pro:mCherry-RCl2A* confocal images after 48h of growth at 21°C or 28°C. (c) Percentage of cells expressing *MUTEpro* after exposure to 21°C or 28°C for 48h and 96h. (d) Confocal in vivo tracing of *MUTEpro:GFP* expression in 3 days-old abaxial cotyledons (0 hours) exposed to 21°C or 28°C during 48h and 96h. Cell contours (purple) show the plasma membrane marker *AtML1pro:mCherry-RCl2A.* Scale bar = 25 µm. (e) Percentage of stomatal fate commitment in precursors expressing *MUTEpro:GFP.* 40 lineages per plant and treatment were traced in 3 different plants (*n* = 3). ns: not significant differences according to Student’s t-test. Bars indicate SEM.

To test if lack of MUTE function increases the meristemoid propensity to become DPs at warm-T, we used *iMUTEmute*, a *mute-3* loss of function line that can be conditionally complemented, easily producing uniform populations of homozygous *mute*-3 seeds (Triviño *et al*., 2013). In non-induced *iMUTEmute* plants, DP proportions after 96h or 11 days at warm-T (Fig. **S5**), did not differ from Col-0 (Fig. **1**) indicating that loss of MUTE function does not promote DP fate.

### Diverted precursors maintain stomatal lineage features

Meristemoid-to-GMC transition has been associated to meristemoid size (Gong *et al.,* 2023). As DPs could derive from a meristemoid subpopulation whose size differs from those progressing to stomata, we traced back meristemoids and DPs (classified by their fate later during lineage development) measuring cell areas at 0h, 48h, 96h and 18 days after transfer to 28°C (Fig. **4a**). The average cell area of stomata-fated meristemoids did not change over the 96h supervised. At 0h, meristemoids with DP or stoma fates had the same size, but after 48h at warm-T, DPs were six-times larger than meristemoids at the same timepoints, and even larger than the few meristemoids present in expanded cotyledons 18 days later.

**Fig. 4.**
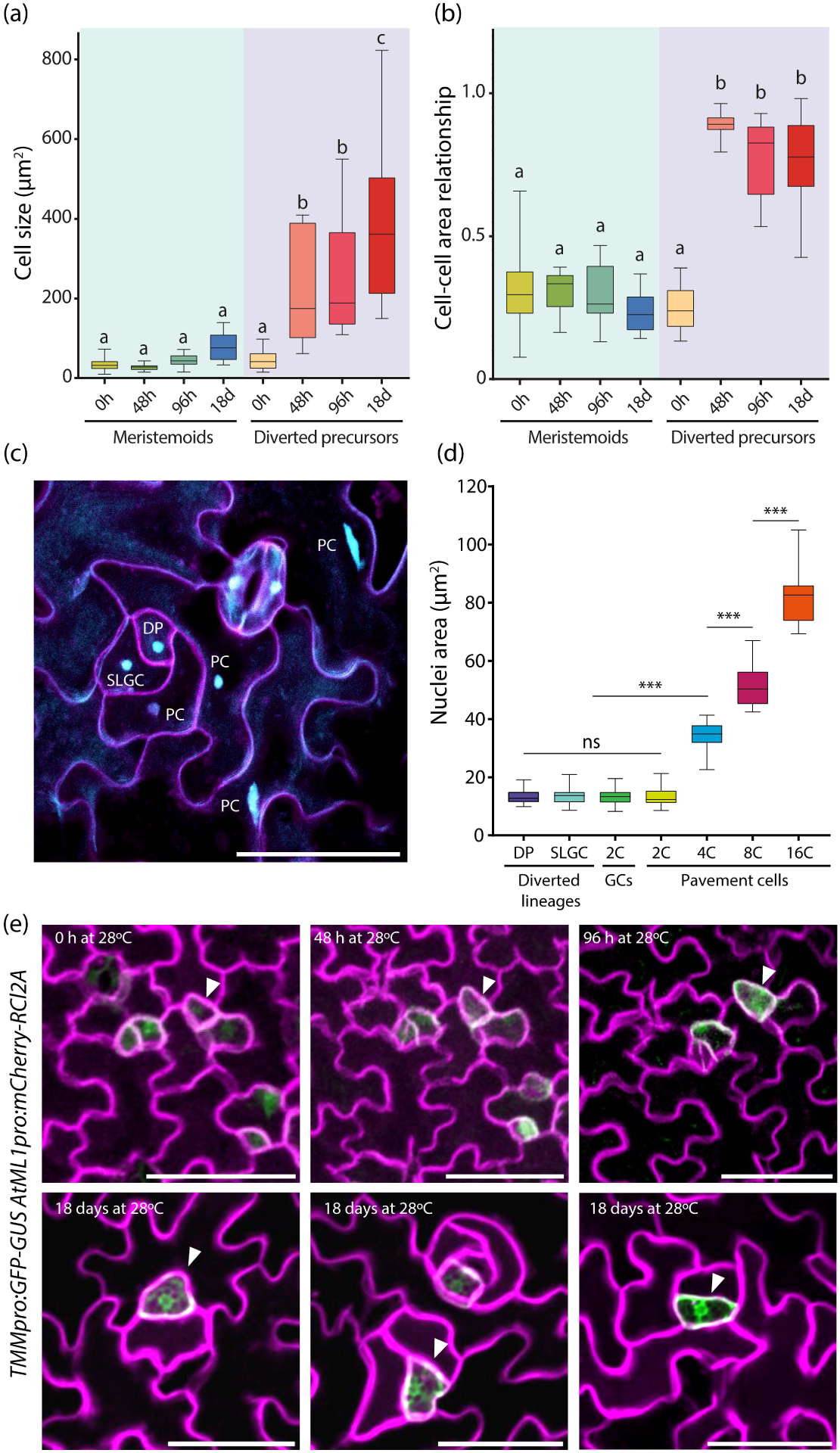
Characterization of diverted precursors. (a,b) Lineages were traced in abaxial cotyledons of the *AtML1pro:mCherry pSPCH:GFP* line in 3-days-old plants transferred to 28°C, and cell sizes determined at 0h, 48h and 96h. Data for the 18 days treatment was obtained from DIC images. (a) Cell size as area of meristemoids and diverted precursors. (b) Area ratio of meristemoid-sister SLGC and DP-sister SLGC pairs. Letters indicate statistical differences according to one-way ANOVA with post hoc Tukey HSD (0h, *n* = 120; 48h, *n* = 38; 96h *n* = 40; 18d *n* = 29 from *n* ≥ 3 independent plants). (c, d) Ploidy level estimated in *AtML1pro:mCherry* plants after 18 days at 28°C. (c) DAPI nuclei staining showing diverted precursors (DP), pavement cells (PC) and SLGC. (d) Nuclei area classified by cell type indicating ploidy levels (as in Triviño *et al*., 2013) in DPs (2C, *n* = 27), SLGCs (2C, *n* = 73), GCs (2C, *n* = 61) and PC (2C, *n* = 71; 4C, *n* = 47; 8C *n* = 55; 16C *n* = 16). Cell nuclei (*n* = 350) were measured in 20 individual plants. Asterisks represent *P* < 0.05 (*), *P* < 0.01 (**), *P* < 0.001 (***) and not significant (ns) differences according to Kruskal-Wallis’s test. Error bars represent SEM. Scale bar = 50 µm. (e) Time-course of *TMMpro:GFP-GUS* transcriptional activity in traced lineages from 3 days-old cotyledons (0h) after 48h and 96h (upper panels) and 18 days (lower panels) exposure to 28°C. White arrowheads indicate *TMMpro:GFP-GUS* signal in diverted precursors.

After 18 days at warm-T, DP nuclear size statistically matched that of 2C guard cells (Fig. **4c,d**), indicating that DPs remain diploid as the meristemoids from which they derive, and did not have higher DNA contents indicative of endoreduplication. SLGCs associated to DPs mostly remained diploid (resembling *mute-3* arrested lineages (Triviño *et al*., 2013)). The size of meristemoids or DPs relative to their sister SLGCs remained constant along time (Fig. **4b**), denoting coordinated cell expansion in meristemoid-SLGC and in DP-SLGC pairs. This hinted that though DPs lost stomatal fate, they maintain characteristics of stomatal lineage cells. In accordance, DPs retained the lineage-specific *TMM* expression (Fig. **4e**). *TMMpro:GFP-GUS* plants grown at warm-T expressed GFP in abundant lineage cells at early developmental stages, contrary to *SPCHpro* and *MUTEpro* silencing. Lineage tracing demonstrated that *TMMpro*-expressing DPs at 96h derive from earlier meristemoids. DPs maintain *TMMpro* expression as late as 18 days after transfer to 28°C, while this marker is restricted to post-protodermal lineage cells in earlier developmental stages (Nadeau & Sack, 2002). This *TMMpro* expression confirms that DPs retain stomatal lineage identity.

### Diverted precursors exit the cell division cycle

Since DPs did not endoreplicate nor divide, we traced cell cycle stages during meristemoid progression to DPs using the multi-fluorescent *PlaCCI Lti6b-GFP* translational reporter line (Kurup *et al*., 2005; Desvoyes *et al*., 2020; Han *et al*., 2022). CDT1a-eCFP marks cells after mitosis that will progress towards G1 or G0 phases, H3.1-mCherry labels cells in S and G2, and CYCB1;1-YFP cells at late-G2 and mitotic commitment. Plants were grown at 28°C from day 3 (0h), recording markers expression and cell divisions at 48h, 96h and 144h (Fig. **5a**). Same size populations of DPs and stomata were selected at 96h, and their histories traced back to 0h and forward to 144h. In lineages that produced a stoma, markers matched the expected behavior (Zuch *et al*., 2023): at 48h, most traced cells had lost CDT1a, showed the enhanced H3.1 signal indicative of S-phase entry (Fig. **5a,b**) and expressed the mitotic commitment marker CYCB1;1. In contrast, a significant proportion of DP-fated cells persistently accumulated CDT1a at 48h and 96h (Fig. **5a,c**), and a few still did after 6 days (114h). H3.1 did not accumulate in DP-fated cells after CDT1a-CFP degradation, indicating that they never entered the S-phase and were therefore stalled in G1 or had exited the cell division cycle towards G0. Then we questioned how differential CDT1a expression related to ACDs in meristemoids with stomata and DP fate. Most ACDs took place during the initial 48h, and their number (around 60% of the traced lineages) did not significantly differed between DP- and stoma-fated meristemoids (Fig. **5a,d**), indicating that during these 48h at 28°C many DP-producing meristemoids retained cell division capacity. However, many DPs expressed CDT1a but never divided, suggesting their cell cycle exit to G0. These were 16% in the 0h to 48h interval, and as many as 40% at later times. All stomata-producing meristemoids that expressed CDT1a experienced a subsequent cell division (Fig. **5e**). Thus, DP-fated meristemoids can divide during the first 48h at warm-T but later they exit the cell cycle and remain in G0.

**Fig. 5.**
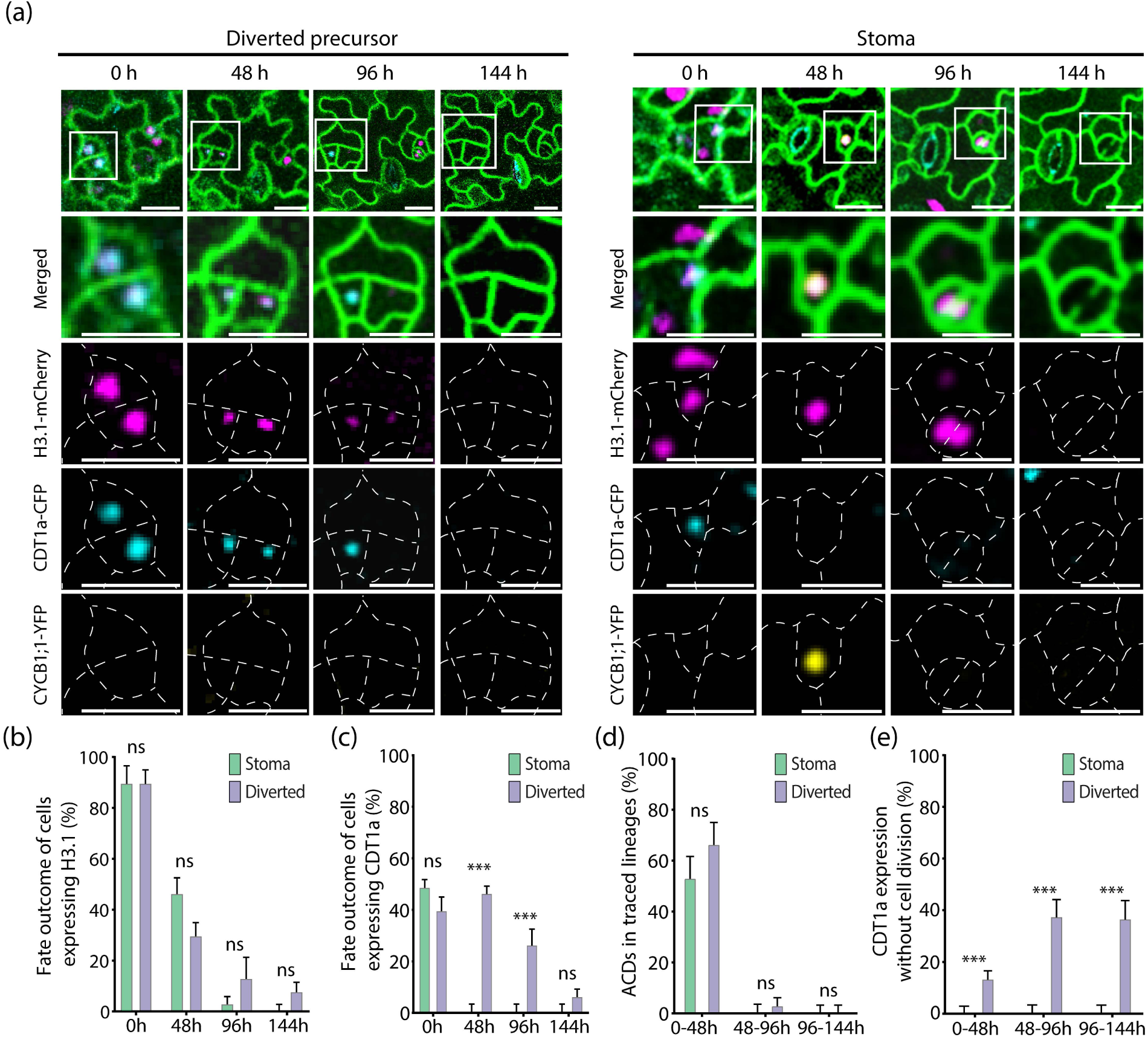
Diverted precursors show altered cell cycle progression. Tracing of lineages resulting in diverted precursor or stoma after exposure to 28°C from day 3 (0 h) to 9 (144h) in the *35S:Lti6b-GFP PlaCCI* multifluorescent reporter line. Cell contours (green) show *35S:Lti6b-GFP*, and nuclei show *H3.1-mCherry* (purple), *CDT1a-CFP* (blue) or *CYCB1;1-YFP* (yellow) expression. Traced lineages were classified according to their fates as stoma or diverted precursor at day 9 (144h). (a) Representative confocal images of traced lineages recording marker expression in the different channels and the merged image at the top. (b,c) Percentage (%) of fate outcome in cells expressing *H3.1-mCherry* or *CDT1a-CFP*. (d) Percentage of ACDs in traced lineages and number of cells expressing *CDT1a* without cell division at 48h intervals. Asterisks represent *P* < 0.05 (*), *P* < 0.01 (**), *P* < 0.001 (***) and not significant (ns) differences according to pairwise Student’s t-test comparison. 3 independent plants were examined, tracing 10 DPs and 10 stomata per plant. Scale bar = 20 µm.

### *SPCHpro* repression induced by warm temperature is reversible

We questioned whether the *SPCHpro* repression triggered by warm-T was reversible. After 16h at 28°C, only 6% of epidermal cells expressed *SPCHpro:GFP*, compared with 15% at control-T (Fig**. 6a,b**). Over the following 8h, normal lineage progression depleted the meristemoid population, and plants kept at 21°C or 28°C lowered the proportion of *SPCHpro*-expressing cells to 10% and 3,7%, respectively. Interestingly, plants grown at 28°C for 16h (T0) and shifted back to 21°C for the following 8h (T1, recovery treatment) increased the proportion of *SPCHpro*-expressing cells to 7,5% (Fig. **6a,b**). We traced individual meristemoids during the recovery period, along with the control at 21°C (Fig. **6c**). Traced meristemoids were classified in four categories of *SPCHpro* behavior during these 8h (Fig. **6c,d**): expressing at T0 and T1 (maintaining expression); expressing at T0 but non-expressing at T1 (loosing expression); non-expressing at T0 nor at T1 (never expressing); non-expressing at T0 and expressing at T1 (gaining expression). In the control-T sample, 80% of the traced meristemoids expressed *SPCHpro* at T0. Due to lineage progression, some had lost expression by T1, but most were still expressing, and a small fraction (10%) gained *SPCHpro* expression. In the recovery treatment, only 30% of the traced meristemoids expressed *SPCHpro* at T0, after 16h at 28°C. At T1 some lost expression as in the control sample but, contrary to the control, the proportion of expressing meristemoids increased to 50%. Of those, around 20% came from meristemoids expressing *SPCHpro* at T0 (maintained expression), but 30% gained expression as they derived from meristemoids non-expressing at T0. Thus, *SPCHpro* repression by warm-T is reversible upon shifting plants to 21°C. Given that DPs cannot be identified at these early times, we could not associate *SPCHpro* behavior with DP fate during the recovery (see Fig. 8 for assessing reversibility of DP fate).

**Fig. 6.**
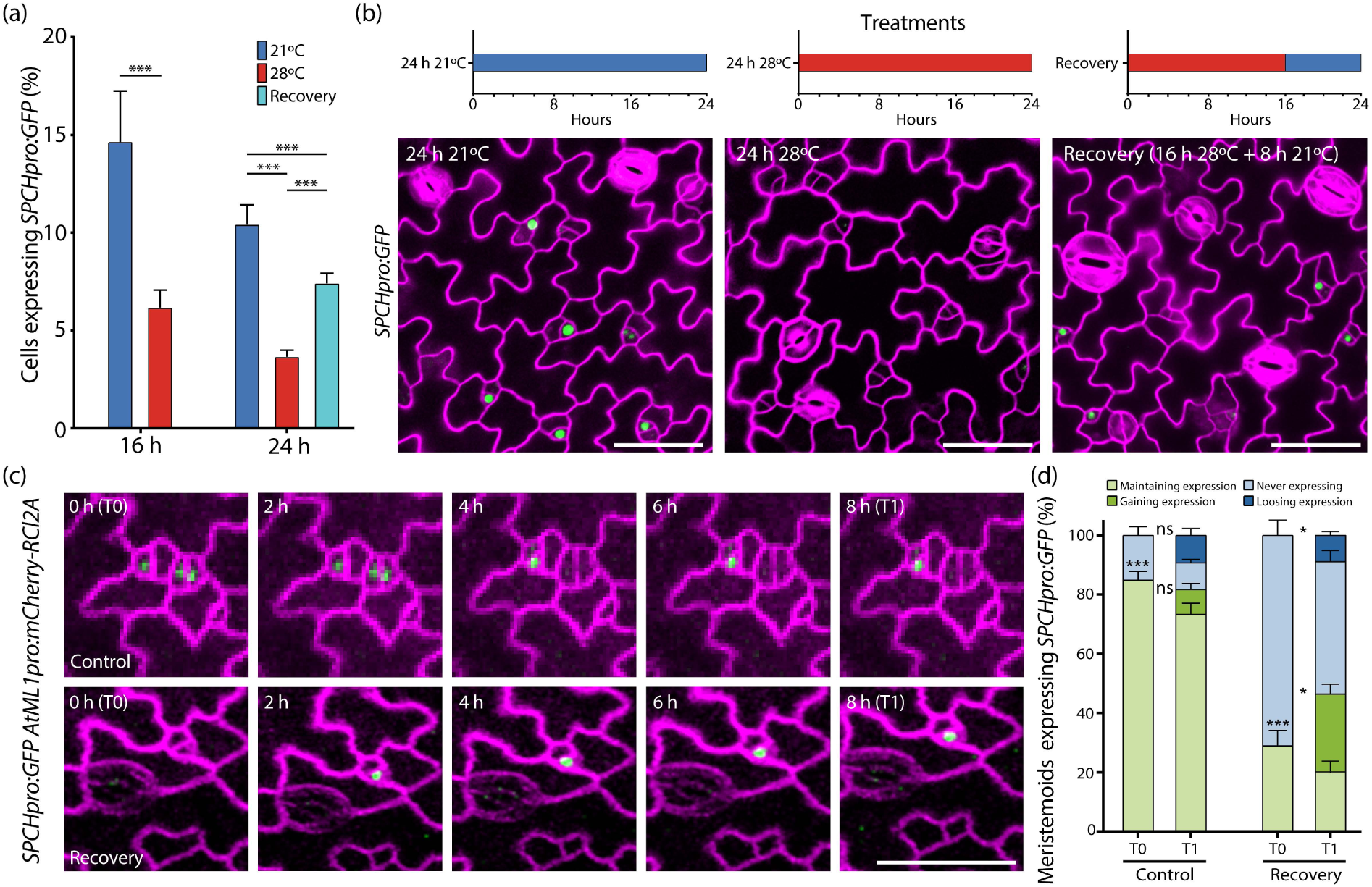
*SPCH* repression by warm temperature is reversible. *SPCHpro:GFP* transcriptional activity in 3 days-old plants grown for 24h at 21°C or 28°C, or kept 16h at 28°C followed by 8h at 21°C (Recovery). (a) Percentage of cells expressing *SPCHpro:GFP* in each treatment. (b) Representative confocal images at the end of each treatment showing GFP (green) and cell contours (purple). (c) Live confocal tracing of representative lineages during the last 8h of control (21°C) and recovery (28°C to 21°C) treatments. (d) Percentage of meristemoids expressing *SPCHpro:GFP* at T0 (16h) and T1 (8h later) during control (21°C) and recovery (28°C to 21°C) treatments, classified as expressing at T0 and T1; expressing at T0 but non-expressing at T1; non-expressing at T0 nor at T1; non-expressing at T0 and expressing at T1. Asterisks represent *P* < 0.05 (*), *P* < 0.01 (**), *P* < 0.001 (***) and not significant (ns) differences according to pairwise Student’s t-test comparison (*n* = 5 plants per treatment). GFP signal was scored in 5 plants per treatment, following 40 meristemoids per plant and treatment (*n* = 200). Scale bar = 25 µm.

### Gene expression changes antecede temperature-induced cell fate decisions

Transcriptional changes had to precede cell fate decisions in stomatal lineages during warm-T and recovery treatments. Lineage cell-type indexes after 16h or 24h at warm or control-T, or in the recovery treatment showed no differences (Fig. **S6**), whereas *SPCHpro* expression experienced significative changes (Fig. **6a**). To analyze global changes in gene expression during these 24h treatments, we first determined *SPCH* and *PIF4* transcript levels by RT-qPCR (Fig. **7a-b**), at 0h, 16h (at 21°C or 28°C) and at 24h (at 21°C, 28°C and in the recovery treatment). As we found diagnostic expression changes, we chose these treatments (as in Fig. 6) for RNAseq. Since at 96h more than 85% of all lineages had formed stomata (Fig. **2**), these lineages are more prominent in our samples than those leading to DPs (8%). RNAseq analysis identified a suit of known stomatal development genes that were differentially expressed (DE) at 16 and 24h between the 21°C and 28°C samples (Fig. **7c**; Table **S2**). *SPCH* transcripts decreased in response to warm-T and among the more down-regulated stomatal genes were several SPCH targets (Lau *et al*., 2014). At 24h, the most temperature-repressed gene was *MUTE*. The recovery treatment largely reverted warm-T induced changes, strongly up-regulating *MUTE* and, to a lesser extent, *SPCH* and other stomatal genes. *PIF4* expression, increased by warm-T, dropped in the recovery.

**Fig. 7.**
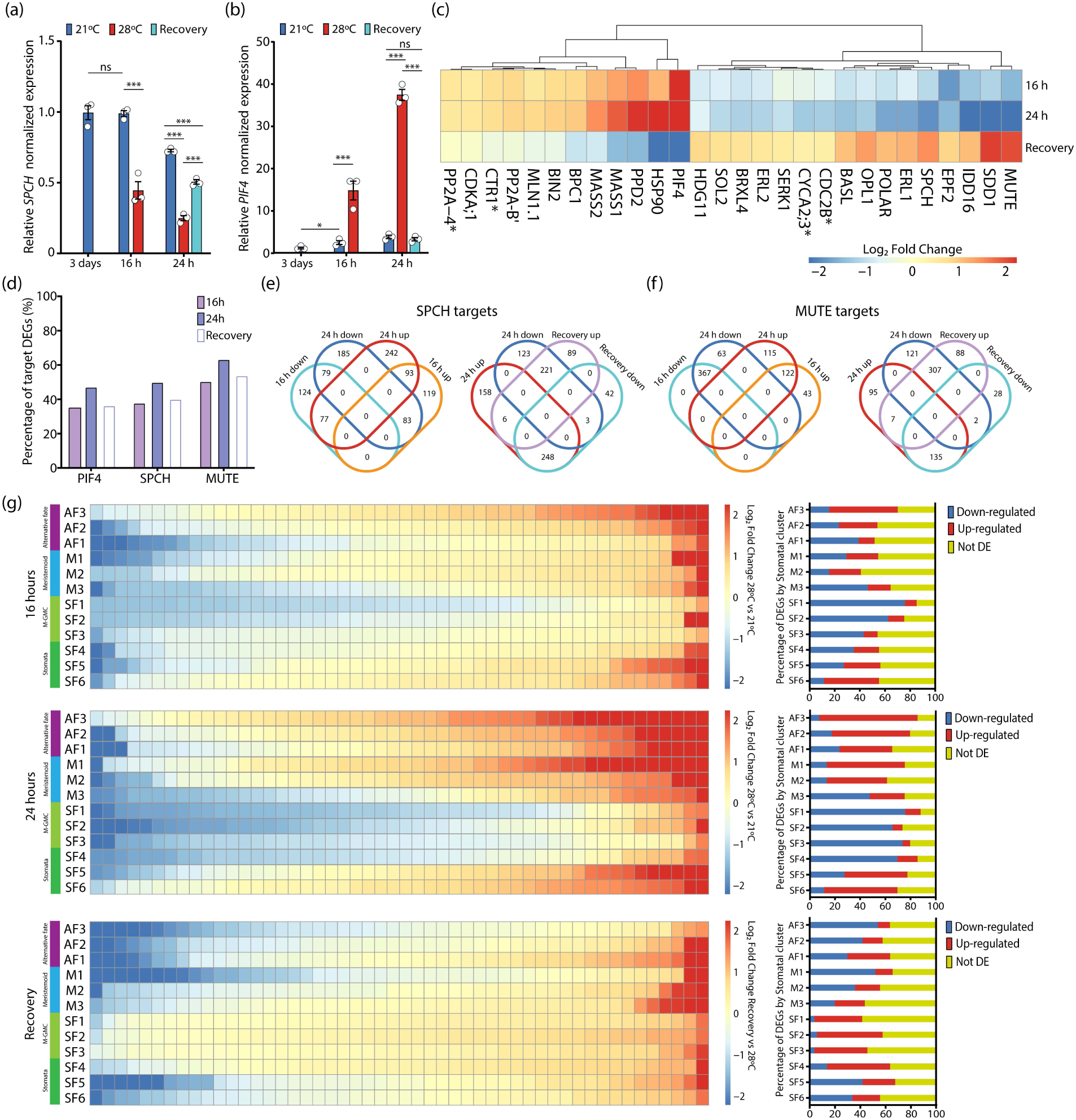
Transcriptional responses to temperature treatments of stomatal lineage genes. (a-b) RT-qPCR analyses of *SPCH* (a) and *PIF4* (b) expression at the indicated time-points for 21°C, 28°C and recovery treatments as in Fig. 6. Expression levels were normalized to *UBIQUITIN10* and *ACTIN2* and expressed relative to levels at the beginning of treatments (3-days-old seedlings = 0h, at 21°C). Error bars represent SEM from three biological replicates. Asterisks represent *P* < 0.05 (*), *P* < 0.01 (**), *P* < 0.001 (***) and non-significant (ns) differences according to pairwise Student’s t-test comparison. (c-g) Transcriptomic analysis by RNAseq of the warm-T response in stomatal lineage genes. (c) Heatmap depicting expression profiles of the stomatal development genes differentially expressed by warm-T at both 16h and 24h or in the recovery treatment. Heatmap values represent change of expression (Log_2_FC) by warm-T (28°C vs. 21°C) at 16h or 24h, and by the recovery treatment (recovery at 24h vs 28°C at 24h). Asterisk (*) represent non-DEGs in the recovery treatment. The complete list of analyzed genes is in Table S1. (d) Percentages of PIF4, SPCH and MUTE targets differentially expressed among treatments. (e-f) Venn diagrams comparing SPCH (e) or MUTE (f) targets differentially expressed among treatments, showing DEGs at 16 and 24h (left), and DEGs at 24h and the recovery treatment (right). (c-f) Genes with FDR < 0.05 and Log_2_FC > |0.5| were considered as differentially expressed. (g) Transcriptional responses to temperature treatments of gene clusters characteristic of distinct cell stages during stomatal lineage development trajectories (described in López-Anido *et al*., 2021). Left panels show heatmaps of expression changes (Log_2_FC) for the top 50 genes per cluster. No Log_2_FC or FDR cut-off were applied. Right panels show percentages of differentially expressed genes (FDR < 0.05) at each time-point and temperature treatment in each gene cluster. Lineage trajectories are meristemoid (m): m1-m3; stomatal fate (sf): sf1-sf6; alternative fate (af): af1-af3.

As SPCH, MUTE and PIF4 are transcription factors, we inspected the behavior of their targets (Lau *et al*., 2014; Pfeiffer *et al*., 2014; Han *et al*., 2018, 2022). About 50% of SPCH and PIF4 targets, and more than 60% of MUTE targets, were DE after 24h at 28°C (Fig. **7d**; Tables **S3** and **S4**). During the recovery, over 60% of MUTE and SPCH targets reverted their temperature-triggered deregulation (Fig. **7e-f**). Transcriptional recovery was more pronounced among down-regulated MUTE targets (71%) including genes involved in the GMC division, and consistent between up- and down-regulated SPCH targets (around 60%).

MUTE targets *FAMA* and *FLP* did not parallel the *MUTE* response to warm-T (Tables **S2** and **S4**). *FAMA* transcripts remained unaffected, and *FLP* transcripts decreased only slightly. However, *FLP,* as *MUTE,* was notably induced during the recovery. Since FLP positively regulates the M-to-GMC transition under compromised MUTE function (Li *et al*., 2023), the sustained *FLP* expression at warm-T might be accountable for most lineages forming stomata despite the deep *MUTE* repression. Analysis of a *FLPpro:GUS-GFP* line showed that the percentage of *FLPpro*-expressing epidermal cells was unaffected by temperature (Fig. **S7**). However, the loss-of-function *flp-1* mutant showed at 28°C lower stomatal unit index and higher DP index than Col-0. Even more, at 21°C, *flp1* produces DPs and has a lower stomatal unit index than Col-0 (Fig. **S8**). These findings support a positive FLP role in stomata fate under warm-T.

To determine if these changes align with transcriptional reprogramming of lineage cell types, we compared our transcriptomes with the transcriptional signatures of distinct cell states and identities from single-cell RNA sequencing (Lopez-Anido *et al*., 2021; Kim *et al*., 2023) (Fig. **7g**; Table **S5;** Fig. **S9**). After 16h at warm-T, genes associated with late-M, M-GMC and early-GMC were consistently down-regulated, while genes associated with stomata differentiation were mostly up-regulated, matching the observed faster lineage progression under warm-T. Notably, alternative fate trajectory genes (Fig. **7g**; Table **S5**) were also up-regulated. All these changes were more pronounced at 24h. The recovery treatment decreased the proportion of up-regulated genes and increased those down-regulated in uncommitted cell states. At the same time, committed cell states had many more up-regulated genes, particularly those from late-M to early-GCs. The undefined stages were able to transition to committed stomatal fates when temperature was lowered. Therefore, warm-T disrupts canonical cell-identities promoting more undefined, flexible cell-identities in the stomatal lineages that had extremely low SPCH and MUTE levels. In addition, lineage progression may rely partly in alternative routes (e.g., involving FLP).

### Developmental window for the reversible stomatal response to warm temperature

Stomata production is established early during organ development, and stomatal lineages initiated later contribute little to final SI. DP fate, paralleled by *SPCHpro* expression reduction, can be acquired after 48 hours at 28°C (Fig. **2**) and *SPCHpro* repression by 16h at warm-T is partly recovered upon shifting plants to control-T (Fig. **6**). To examine whether DP-fate is also reversible and how the length of exposure to warm-T impacted DPI and SI in mature organs, seedlings were exposed to warm-T for increasing times (16h as in the recovery, 24h, 48h, 96h, and 144h; Fig. **8a**) before shifting them back to 21°C or keeping them at 28°C, determining SI and DPI in the mature cotyledon, at 21 days.

**Fig. 8.**
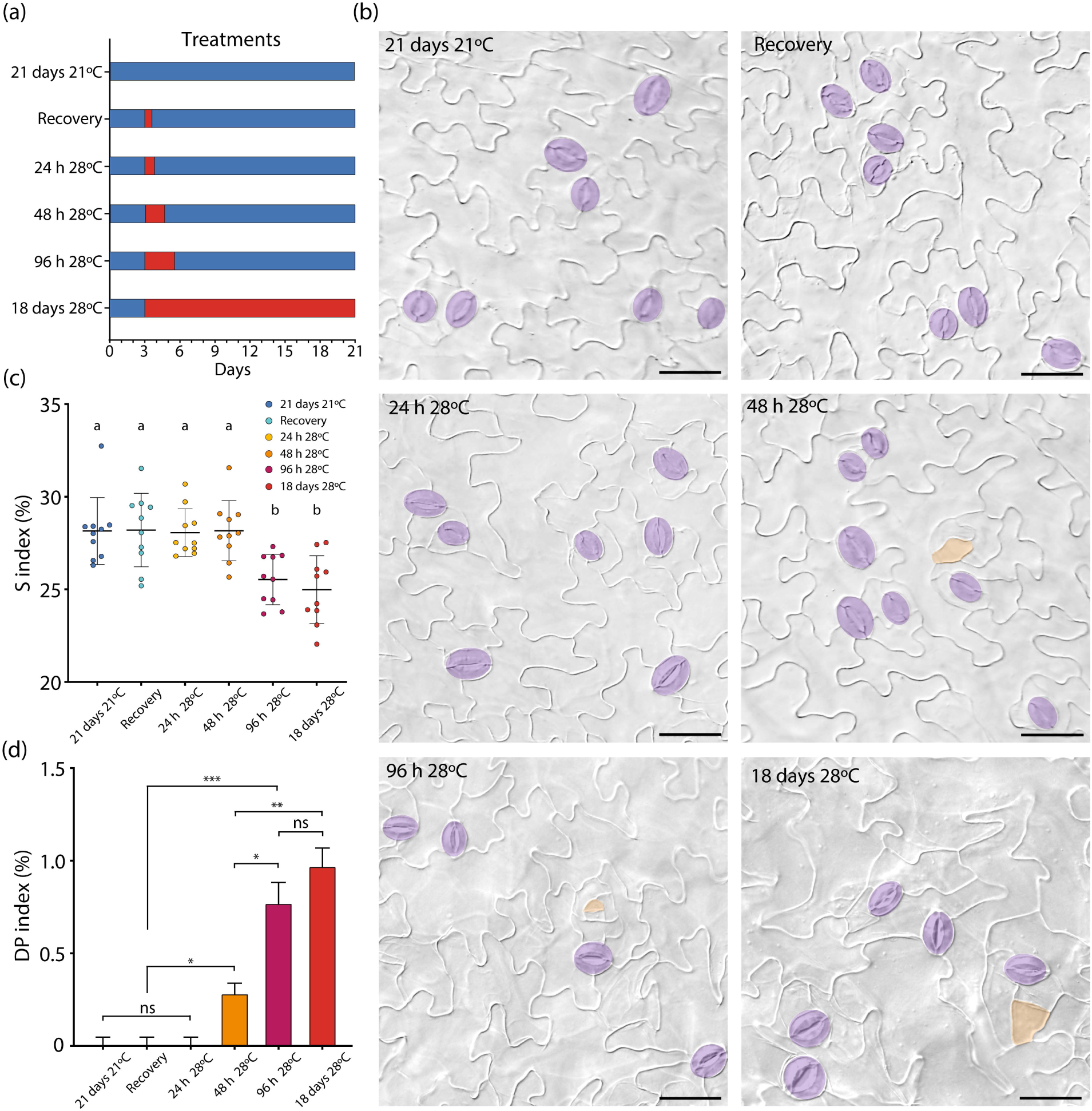
Irreversible stomatal index reduction and diverted precursor formation requires a minimum time length of warm-T exposure. (a) Schematic representation of temperature treatments. 3-days-old Col-0 seedlings were exposed to increasingly longer warm-T treatments (0h, 16h -recovery-, 24h, 48h, 96h or 21 days) before shifted back to 21°C and monitored at 21 days. (b) Representative DIC micrographs of abaxial mature cotyledons at 21 days after the treatments. Scale bar = 50 µm. Stomata are colored in purple and diverted precursors in orange. (c) Stomatal index of mature 21-days-old cotyledons at the indicated treatments. Plots show mean ± SD. Each point represents the value of an individual plant. Letters indicate statistical differences according to one-way ANOVA with post hoc Tukey HSD. (d) Diverted precursor index in the same samples. Asterisks in (b) and (c) represent *P* < 0.05 (*), *P* < 0.01 (**), *P* < 0.001 (***) and not significant (ns) differences according to pairwise Student’s t-test comparison. (c-d) *n* = 10 plants per treatment. Bars represent SEM.

Only plants grown for at least 96 hours at 28°C showed the same SI reduction than those continuously grown at 28°C (Fig. **8b,c**) and had a similar DP production (Fig. **8d**). However, plants grown at 28°C for only 48h produced DPs, although this did not translate to SI reduction (Fig **8c,d**). Shorter exposure to warm-T did not produce DPs nor SI decreases. Thus, DP identity is irreversibly acquired after exposition to warm-T for at least 48h, and DP production determines the SI index reduction only if warm-T is maintained at least 96 hours.

The modest (around 3%) SI decrease triggered by warm-T or PIF4-overexpression suggests a limited physiological impact of this developmental response. But the trait with physiological consequences is stomatal density (SD), which in induced *iPIF4oe* and warm-T-treated Col-0 plants had values *c.* 30% lower than plants at control-T (Fig. **S10**). This notable SD reduction accompanying the small SI changes would have a strong impact on plant physiology and suggests a relevant adaptive nature of the T-induced SI decrease.

## DISCUSSION

### Loss of stomatal precursor identity underly SI reduction induced by warm-temperature or PIF4-overexpression

The SI reduction induced by warm-T was ascribed to a negative transcriptional feedback loop between *SPCH* and *PIF4* (Lau *et al*., 2018). However, the underlying developmental basis of this mechanism was unaccounted for. We discovered that warm-T or *PIF4*-overexpression induce diverted stomatal lineage precursors (DPs) unable to form stomata, whose prevalence consistently account for the SI reduction observed (Fig. **1**). *PIF4* overexpression produced DPs in a temperature-independent manner and, remarkably, *iPIFoe* growth at 28°C did not increase DP numbers further, indicating a saturated response. Higher *SPCH* levels (Fig. **S4**) counteracted warm-T-dependent DP formation and SI reduction, while inducing a small DP proportion at control-T, what indicates that higher than normal *SPCH* levels can disrupt meristemoid fates. Our findings fit with previous results (Lopez-Anido *et al*., 2021), describing diverted cells induced by impaired *SPCH* transcription, morphologically similar to DPs.

We demonstrate that DPs derive from meristemoids by following *SPCHpro* activity and tracing individual lineage histories early during development at 28°C. Since DPs always derive from meristemoids that lost *SPCH* expression, *SPCH* silencing is required for DP fate. DP-fated meristemoids acquired larger sizes and the DP/SLGC size ratio, close to 0.3 immediately after the ACD as in normal meristemoid/SLGC pairs, increased to almost 0.8 after 48h. Coordinated expansion of DP/SLGC fits with the need of late-meristemoid progression for SLGC expansion (Triviño *et al*., 2013). DPs remained diploid and expressed *TMMpro* and, after 48h at warm-T, half of them accumulated CDT1a without subsequent cell division, and when CDT1a extinguished, this was not followed by nuclear H3.1 incorporation indicative of S-phase. These findings imply that DPs enter in G0. Their cell-cycle exit could be forced by very low SPCH and MUTE levels that compromise cell divisions. Thus, DPs differ from *mute* meristemoids, which retain *SPCH* expression and where G1 is progressively extended in repeated ACDs (Zuch *et al*., 2023). in DPs, SPCH is almost absent, and they rarely divide (Fig. **2a**). All these features indicate that DPs are unique lineage cells that lost stomatal fate and division potential.

### DP fate acquisition precedes *MUTE* expression

The late-meristemoid transition to GMC, committed to form a stoma, requires MUTE (Han *et al*., 2018). Tracing *MUTEpro:GFP* we found that warm-T drastically reduces *MUTEpro*-expressing cells, and that DPs derive only from precursors that did not express *MUTE*. All lineages that expressed *MUTE* formed stomata, indicating that *MUTE* expression makes late-meristemoids insensitive to warm-T. The absence of MUTE function does not promote DP fate, as in the loss-of-function *mute-3*, warm-T induces the same DP proportion observed in Col-0.

### Stomatal lineages progress at warm-T through rewired transcriptional networks

DP proportions are small, at warm-T or in *iPIF4oe*. It is striking that, despite *SPCH* silencing, most lineages formed stomata. The reciprocal PIF4/SPCH loop -where *SPCH* repression by PIF4 is counteracted by *PIF4* repression by SPCH-was presumed to ensure enough SPCH to form abundant stomata at warm-T (Lau *et al*., 2018). However, in *iPIF4oe* this loop does not operate, as PIF4 is under a constitutive promoter not repressed by SPCH. It could be argued that in cotyledons the 3-days of 21°C exposure primes SPCH function prior to warm-T, but this does not apply to *iPIFoe* nor to first leaves, which produce abundant stomata at 28°C. *SPCH* transcript levels are extremely low but not zero, perhaps sufficient to impulse most lineages, but this would not compensate the even deeper drop in *MUTE* expression. Therefore, alternative developmental pathways possibly drive lineage progression and stomatal fate commitment. Indeed, we found that warm-T accelerates stomata formation (Fig. **2d**). Faster development at warm-T, including leaf growth, has been established (Casal & Balasubramanian, 2019) and several stomatal mutants also exhibited it (Pérez-Bueno *et al*., 2022), but how stomatal development is altered has not been reported. While warm-T accelerates cell division in root meristems (Ai *et al*., 2023), faster growth in aerial organs was ascribed to increased cell expansion. The accelerated lineage development that we observed might relate to changes in ACD and SCD length, estimated at approximately 12h and 20h, respectively, under control-T (Han *et al*., 2022). Consequently, 48 hours at warm-T would suffice for several ACDs and the terminal SCD.

During these critical 48h at warm-T, *SPCHpro* and *MUTEpro* are silenced, diploid *TMMpro*-expressing DPs are specified and enter G0, but most lineages culminate their development and form stomata. Live lineage tracing showed *SPCHpro* silencing in the first 16h that increased 8h later (Fig. **6**). But if, after 16h at 28°C, plants are shifted to 21°C, *SPCHpro* expression is partially recovered. The increase in *SPCHpro*-expressing cells came partly from previously expressing meristemoids but partly from silenced meristemoids, demonstrating that *SPCHpro*-repression triggered by a warm-T pulse is reversible. Murcia *et al*. (2022) reported that the increase in *PIF4* promoter activity triggered by warm-T during the day was irreversible during cooler nights. This contrasts with our finding that *PIF4* transcript accumulation during the 28°C exposure in daylight was drastically reduced following an 8-hour recovery at 21°C in darkness. Differences in photoperiod (long-day versus short-day), night temperature (21°C in our experiments and 10°C in theirs) and light quality may account for the contrasting results.

The disparity in stomatal precursor behaviors among these 24h-long treatments predicted distinct lineage-associated transcriptional landscapes. Because more than 80% of lineages at either temperature produce stomata, transcriptional differences would mostly represent these stomata-producing lineages. Warm-T produced profound and consistent changes in the expression of stomatal development genes, most recovered after 8h at 21°C (Table **S2**, Fig **7**). Many of the warm-T triggered expression changes fit with *SPCHpro* and *MUTEpro* silencing and presumably contribute to reducing SPCH and MUTE activity, ACDs, and stomata-producing SCDs, raising the question of how most lineages form stomata under these conditions. Although most MUTE transcriptional targets are DE in opposite directions by warm-T and by the recovery treatment, some were not. Among MUTE targets not repressed by warm-T was *FLP*. Since FLP can sustain the M-to-GMC transit under impaired MUTE function (Li *et al*., 2023) it may contribute to lineage completion at warm-T. This is supported by our finding that *flp-1* mutants formed more DPs than Col-0 at 21°C and 28°C (Fig. **S8**). Thus, the severe *MUTE* silencing could be overcome through unconventional pathways, such as that mediated by FLP, driving stomatal fate in most lineages.

Regarding lineage progression, while *SPCH* was strongly -though not completely-repressed, warm-T promoted expression changes predicted to increase SPCH stability. *MAPK SUBSTRATES IN THE STOMATAL LINEAGE 1/2* (*MASS1/2*), positive drivers of stomatal development that increase SPCH stability interfering with the YODA-led repressive MAPK cascade, were strongly up-regulated (Qi *et al*., 2019; Xue *et al*., 2020). Warm-T decreased transcripts for several negative regulators (for instance, *SDD1*, *ER*, *ERL1*/*2* and *EPF2*), potentially relieving repression of SPCH (and MUTE) activities. In this sense, SPCH requirement can be overcome by ectopic *MUTE* expression (Pillitteri *et al*., 2007), and stomata can be made in the absence of MUTE if negative regulators, as TMM or ERf, are missing (Pillitteri *et al*., 2008). It is plausible that at warm-T, low *SPCH* expression facilitates lineage progression through increased protein stability. In our experiments, residual SPCH activity generated prior to the exposure to warm-T might also promote SPCH-dependent lineage stages, supported by normal *SCRM/2* levels, not affected by warm-T.

All these results indicate a transcriptionally non-canonical lineage progression at warm-T. Gene clusters defining distinct cell identities (Kim *et al*., 2023) and cell stages during stomatal lineage trajectories (Lopez-Anido *et al*., 2021) showed expression differences between 21°C and 28°C datasets (Fig. **S9** and Fig. **7g**; Table **S4**). Our RNAseq data reveal that warm-T severely constrains identities/stages related to stomata-fate commitment (late-meristemoid, GMC and early GCs), while promoting either alternative-fates or final stoma differentiation through up-regulation of their associated genes. The recovery treatment corrects these trends, decreasing alternative fate-characteristic gene expression and promoting expression of genes associated to stomatal commitment, SCD and early GC differentiation dominated by *MUTE*. Consequently, most lineages form stomata at warm-T, but through reorganized gene networks. The predominant uncommitted stages perhaps favor DP formation that results in SI reduction. We identified known stomatal development genes that should potentially contribute to lineage progression, as *FLP*, but the rewired gene circuits could also use new players that were not identified in control conditions.

### Final remarks

Our work has limitations imposed by the nature of lineage development. Because lineages progress faster at 28°C, most have formed stomata by 48h (Fig. **2b**), precluding molecular analysis of developing lineages later than 24h. Circadian rhythm effects on transcription (Maric & Mas, 2020; Swift *et al*., 2022; Laosuntisuk & Doherty, 2022) are over imposed to our treatments.

SI reductions promoted by warm-T or *PIF4*-overexpression are modest, but are accompanied by much larger decreases in stomatal densities (Fig. **S10**), the trait that influences transpiration and photosynthesis and thus refrigeration and water use efficiency (Franks *et al*., 2015; Harrison *et al*., 2020). The occurrence of distinct inter- and intraspecific stomatal responses to warm-T (Jumrani *et al*., 2017; Caine *et al*., 2019) adds to the well-established role of other heat-adaptive behaviors (Zhu *et al*., 2021), further supporting its relevance under a warming climate. Short-term temperature oscillations prevail in natural conditions, as days are warmer than nights. The need of a longer than 48h exposure to 28°C (Fig. **8**) to trigger DPs that we discovered would prevent an irreversible SI reduction after occasional temperature rises, while promoting uncommitted lineage stages provides developmental flexibility under these conditions.

## Acknowledgments

This work was supported by grants of the Spanish (PID2019-105362RB-I00 and PID2022-137606NB-I00), and the Castilla-La Mancha Governments (SBPLY/21/180225/000058) to MM and CF. The laboratory received support by UCLM intramural grants and EU FEDER funds. JSP received an exchange grant from EMBO. We thank Ana Rapp for technical support.

## Author contributions

JSP performed the research. JSP and MM designed the research. JSP, JIM and BG developed the lines. JSP, AB, LS and MV analyzed RNAseq data. EK and ER helped with confocal imaging. AB, BD, CG and ER contributed to data interpretation. JSP, CF and MM wrote the paper with the contributions of all other authors. CF and MM obtained the funding.

## Data availability

RNAseq data are publicly available in the NCBI’s Gene Expression Omnibus under GEO Series accession no. GSE278519.

## Supporting information

### Legends to Figures S1-S10

**Fig. S1. Loss-of-function *pif* mutants do not produce diverted precursors at warm temperature.**

**Fig. S2. *PIF4* overexpression mimics warm temperature at early developmental times.**

**Fig. S3. Leaf diverted precursors derive from meristemoids.**

**Fig. S4. Conditional SPCH overexpression buffers the stomatal response to warm temperature.**

**Fig. S5. Loss of MUTE function does not influence DP formation.**

**Fig. S6. Cell-type indexes remain unchanged in the 16h and 24h samples used for transcript analysis.**

**Fig. S7. FOUR LIPS promoter is expressed similarly across temperature treatments.**

**Fig. S8. Loss of FLP function increases diverted precursor production.**

**Fig. S9. Transcriptional responses to temperature treatments of genes defining cell identities in the stomatal lineage.**

**Fig. S10. Modest warm-T-induced stomatal index changes concur with broader changes in stomatal density.**

### Tables S1-S5

Table S1. Primers used in this study.

Table S2. Transcriptional response of known stomatal development genes to warm temperature treatments.

Table S3. SPCH targets differentially expressed by warm-temperature treatments. Table S4. MUTE targets differentially expressed by warm-temperature treatments.

Table S5. Temperature-induced expression changes for top 50 genes per cluster in Figure 7g.

## References

Ai H, Bellstaedt J, Bartusch KS, Eschen-Lippold L, Babben S, Balcke GU, Tissier A, Hause B, Andersen TG, Delker C, et al. 2023. Auxin-dependent regulation of cell division rates governs root thermomorphogenesis. The EMBO Journal 42: e111926.

Balasubramanian S, Sureshkumar S, Lempe J, Weigel D. 2006. Potent Induction of Arabidopsis thaliana Flowering by Elevated Growth Temperature. PLOS Genetics 2: e106.

Bergmann DC, Sack FD. 2007. Stomatal Development. Annual Review of Plant Biology 58: 163–181.

Bernardo-García S, De Lucas M, Martínez C, Espinosa-Ruiz A, Davière J-M, Prat S. 2014. BR-dependent phosphorylation modulates PIF4 transcriptional activity and shapes diurnal hypocotyl growth. Genes & Development 28: 1681–1694.

Bertolino LT, Caine RS, Gray JE. 2019. Impact of Stomatal Density and Morphology on Water-Use Efficiency in a Changing World. Frontiers in Plant Science 10.

Bolger AM, Lohse M, Usadel B. 2014. Trimmomatic: a flexible trimmer for Illumina sequence data. Bioinformatics 30: 2114–2120.

Caine RS, Yin X, Sloan J, Harrison EL, Mohammed U, Fulton T, Biswal AK, Dionora J, Chater CC, Coe RA, et al. 2019. Rice with reduced stomatal density conserves water and has improved drought tolerance under future climate conditions. The New Phytologist 221: 371–384.

Casal JJ, Balasubramanian S. 2019. Thermomorphogenesis. Annual Review of Plant Biology 70: 321–346.

Clough SJ, Bent AF. 1998. Floral dip: a simplified method for *Agrobacterium*-mediated transformation of *Arabidopsis thaliana*. The Plant Journal 16: 735–743.

Coego A, Brizuela E, Castillejo P, Ruíz S, Koncz C, del Pozo JC, Piñeiro M, Jarillo JA, Paz-Ares J, León J, et al. 2014. The TRANSPLANTA collection of Arabidopsis lines: a resource for functional analysis of transcription factors based on their conditional overexpression. The Plant Journal: For Cell and Molecular Biology 77: 944–953.

Crawford AJ, McLachlan DH, Hetherington AM, Franklin KA. 2012. High temperature exposure increases plant cooling capacity. Current biology: CB 22: R396–397.

Delgado D, Alonso-Blanco C, Fenoll C, Mena M. 2011. Natural variation in stomatal abundance of Arabidopsis thaliana includes cryptic diversity for different developmental processes. Annals of Botany 107: 1247–1258.

Delker C, Quint M, Wigge PA. 2022. Recent advances in understanding thermomorphogenesis signaling. Current Opinion in Plant Biology 68: 102231.

Desvoyes B, Arana-Echarri A, Barea MD, Gutierrez C. 2020. A comprehensive fluorescent sensor for spatiotemporal cell cycle analysis in Arabidopsis. Nature Plants 6: 1330–1334.

Dittberner H, Korte A, Mettler-Altmann T, Weber APM, Monroe G, de Meaux J. 2018. Natural variation in stomata size contributes to the local adaptation of water-use efficiency in Arabidopsis thaliana. Molecular Ecology 27: 4052–4065.

Dow GJ, Berry JA, Bergmann DC. 2014. The physiological importance of developmental mechanisms that enforce proper stomatal spacing in Arabidopsis thaliana. The New Phytologist 201: 1205–1217.

Erwin JE, Heins RD, Karlsson MG. 1989. Thermomorphogenesis in Lilium longiflorum. American Journal of Botany 76: 47–52.

Franklin KA, Lee SH, Patel D, Kumar SV, Spartz AK, Gu C, Ye S, Yu P, Breen G, Cohen JD, et al. 2011. PHYTOCHROME-INTERACTING FACTOR 4 (PIF4) regulates auxin biosynthesis at high temperature. Proceedings of the National Academy of Sciences 108: 20231–20235.

Franks PJ, Drake PL, Beerling DJ. 2009. Plasticity in maximum stomatal conductance constrained by negative correlation between stomatal size and density: an analysis using *Eucalyptus globulus*. Plant, Cell & Environment 32: 1737–1748.

Franks PJ, W. Doheny-Adams T, Britton-Harper ZJ, Gray JE. 2015. Increasing water-use efficiency directly through genetic manipulation of stomatal density. New Phytologist 207: 188–195.

Geisler M, Nadeau J, Sack FD. 2000. Oriented Asymmetric Divisions That Generate the Stomatal Spacing Pattern in Arabidopsis Are Disrupted by the *too many mouths* Mutation. The Plant Cell 12: 2075–2086.

Gong Y, Dale R, Fung HF, Amador GO, Smit ME, Bergmann DC. 2023. A cell size threshold triggers commitment to stomatal fate in Arabidopsis. Science Advances 9: eadf3497.

Gray WM, Östin A, Sandberg G, Romano CP, Estelle M. 1998. High temperature promotes auxin-mediated hypocotyl elongation in Arabidopsis. Proceedings of the National Academy of Sciences 95: 7197–7202.

Hachez C, Ohashi-Ito K, Dong J, Bergmann DC. 2011. Differentiation of Arabidopsis Guard Cells: Analysis of the Networks Incorporating the Basic Helix-Loop-Helix Transcription Factor, FAMA. Plant Physiology 155: 1458–1472.

Han S-K, Qi X, Sugihara K, Dang JH, Endo TA, Miller KL, Kim E, Miura T, Torii KU. 2018. MUTE Directly Orchestrates Cell State Switch and the Single Symmetric Division to Create Stomata. Plant Biology.

Han S-K, Torii KU. 2016. Lineage-specific stem cells, signals and asymmetries during stomatal development. Development 143: 1259–1270.

Han S-K, Yang J, Arakawa M, Iwasaki R, Sakamoto T, Kimura S, Kim E-D, Torii KU. 2022. Deceleration of cell cycle underpins a switch from proliferative- to terminal division in plant stomatal lineage. Plant Biology.

Harrison EL, Arce Cubas L, Gray JE, Hepworth C. 2020. The influence of stomatal morphology and distribution on photosynthetic gas exchange. The Plant Journal 101: 768–779.

Herrmann A, Torii KU. 2021. Shouting out loud: signaling modules in the regulation of stomatal development. Plant Physiology 185: 765–780.

Ishige F, Takaichi M, Foster R, Chua N, Oeda K. 1999. A G-box motif (GCCACGTGCC) tetramer confers high-level constitutive expression in dicot and monocot plants. The Plant Journal 18: 443–448.

Jumrani K, Bhatia VS, Pandey GP. 2017. Impact of elevated temperatures on specific leaf weight, stomatal density, photosynthesis and chlorophyll fluorescence in soybean. Photosynthesis Research 131: 333–350.

Kagan ML, Novoplansky N, Sachs T. 1992. Variable Cell Lineages form the Functional Pea Epidermis. Annals of Botany 69: 303–312.

Kanaoka MM, Pillitteri LJ, Fujii H, Yoshida Y, Bogenschutz NL, Takabayashi J, Zhu J-K, Torii KU. 2008. *SCREAM/ICE1* and *SCREAM2* Specify Three Cell-State Transitional Steps Leading to *Arabidopsis* Stomatal Differentiation. The Plant Cell 20: 1775–1785.

Kim E-J, Zhang C, Guo B, Eekhout T, Houbaert A, Wendrich JR, Vandamme N, Tiwari M, Simon-Vezo C, Vanhoutte I, et al. 2023. Cell type–specific attenuation of brassinosteroid signaling precedes stomatal asymmetric cell division. Proceedings of the National Academy of Sciences 120: e2303758120.

Koini MA, Alvey L, Allen T, Tilley CA, Harberd NP, Whitelam GC, Franklin KA. 2009. High Temperature-Mediated Adaptations in Plant Architecture Require the bHLH Transcription Factor PIF4. Current Biology 19: 408–413.

Kurup S, Runions J, Köhler U, Laplaze L, Hodge S, Haseloff J. 2005. Marking cell lineages in living tissues. The Plant Journal 42: 444–453.

Lai LB, Nadeau JA, Lucas J, Lee E-K, Nakagawa T, Zhao L, Geisler M, Sack FD. 2005. The Arabidopsis R2R3 MYB Proteins FOUR LIPS and MYB88 Restrict Divisions Late in the Stomatal Cell Lineage. The Plant Cell 17: 2754–2767.

Laosuntisuk K, Doherty CJ. 2022. The intersection between circadian and heat-responsive regulatory networks controls plant responses to increasing temperatures. Biochemical Society Transactions 50: 1151–1165.

Lau OS, Davies KA, Chang J, Adrian J, Rowe MH, Ballenger CE, Bergmann DC. 2014. Direct roles of SPEECHLESS in the specification of stomatal self-renewing cells. Science 345: 1605–1609.

Lau OS, Song Z, Zhou Z, Davies KA, Chang J, Yang X, Wang S, Lucyshyn D, Tay IHZ, Wigge PA, et al. 2018. Direct Control of SPEECHLESS by PIF4 in the High-Temperature Response of Stomatal Development. Current Biology 28: 1273–1280.e3.

Lee LR, Bergmann DC. 2019. The plant stomatal lineage at a glance. Journal of Cell Science 132: jcs228551.

Lee E, Lucas JR, Sack FD. 2014. Deep functional redundancy between FAMA and FOUR LIPS in stomatal development. The Plant Journal 78: 555–565.

Li P, Chen L, Gu X, Zhao M, Wang W, Hou S. 2023. FOUR LIPS plays a role in meristemoid- to-GMC fate transition during stomatal development in Arabidopsis. The Plant Journal 114: 424–436.

Lopez-Anido CB, Vatén A, Smoot NK, Sharma N, Guo V, Gong Y, Anleu Gil MX, Weimer AK, Bergmann DC. 2021. Single-cell resolution of lineage trajectories in the Arabidopsis stomatal lineage and developing leaf. Developmental Cell 56: 1043–1055.e4.

MacAlister CA, Ohashi-Ito K, Bergmann DC. 2007. Transcription factor control of asymmetric cell divisions that establish the stomatal lineage. Nature 445: 537–540.

de Marcos A, Houbaert A, Triviño M, Delgado D, Martín-Trillo M, Russinova E, Fenoll C, Mena M. 2017. A Mutation in the bHLH Domain of the SPCH Transcription Factor Uncovers a BR-Dependent Mechanism for Stomatal Development. Plant Physiology 174: 823–842.

Maric A, Mas P. 2020. Chromatin Dynamics and Transcriptional Control of Circadian Rhythms in Arabidopsis. Genes 11: 1170.

McCarthy DJ, Chen Y, Smyth GK. 2012. Differential expression analysis of multifactor RNA-Seq experiments with respect to biological variation. Nucleic Acids Research 40: 4288–4297.

Murcia G, Nieto C, Sellaro R, Prat S, Casal JJ. 2022. Hysteresis in PHYTOCHROME-INTERACTING FACTOR 4 and EARLY-FLOWERING 3 dynamics dominates warm daytime memory in Arabidopsis. The Plant Cell 34: 2188–2204.

Nadeau JA, Sack FD. 2002. Control of Stomatal Distribution on the *Arabidopsis* Leaf Surface. Science 296: 1697–1700.

Ohashi-Ito K, Bergmann DC. 2006. *Arabidopsis* FAMA Controls the Final Proliferation/Differentiation Switch during Stomatal Development. The Plant Cell 18: 2493–2505.

Patro R, Duggal G, Love MI, Irizarry RA, Kingsford C. 2017. Salmon provides fast and bias-aware quantification of transcript expression. Nature Methods 14: 417–419.

Pérez-Bueno ML, Illescas-Miranda J, Martín-Forero AF, de Marcos A, Barón M, Fenoll C, Mena M. 2022. An extremely low stomatal density mutant overcomes cooling limitations at supra-optimal temperature by adjusting stomatal size and leaf thickness. Frontiers in Plant Science 13: 919299.

Pfeiffer A, Shi H, Tepperman JM, Zhang Y, Quail PH. 2014. Combinatorial complexity in a transcriptionally centered signaling hub in Arabidopsis. Molecular Plant 7: 1598–1618.

Pillitteri LJ, Bogenschutz NL, Torii KU. 2008. The bHLH Protein, MUTE, Controls Differentiation of Stomata and the Hydathode Pore in Arabidopsis. Plant and Cell Physiology 49: 934–943.

Pillitteri LJ, Sloan DB, Bogenschutz NL, Torii KU. 2007. Termination of asymmetric cell division and differentiation of stomata. Nature 445: 501–505.

Pillitteri LJ, Torii KU. 2012. Mechanisms of Stomatal Development. Annual Review of Plant Biology 63: 591–614.

Qi S, Lin Q, Feng X, Han H, Liu J, Zhang L, Wu S, Le J, Blumwald E, Hua X. 2019. IDD 16 negatively regulates stomatal initiation via trans-repression of *SPCH* in *Arabidopsis*. Plant Biotechnology Journal 17: 1446–1457.

Qi X, Torii KU. 2018. Hormonal and environmental signals guiding stomatal development. BMC Biology 16: 21.

Quint M, Delker C, Franklin KA, Wigge PA, Halliday KJ, van Zanten M. 2016. Molecular and genetic control of plant thermomorphogenesis. Nature Plants 2: 1–9.

Raven JA. 2002. Selection pressures on stomatal evolution. The New Phytologist 153: 371–386.

Schneider CA, Rasband WS, Eliceiri KW. 2012. NIH Image to ImageJ: 25 years of image analysis. Nature Methods 9: 671–675.

Soneson C, Love MI, Robinson MD. 2016. Differential analyses for RNA-seq: transcript-level estimates improve gene-level inferences. F1000Research 4: 1521.

Swift J, Greenham K, Ecker JR, Coruzzi GM, Robertson McClung C. 2022. The biology of time: dynamic responses of cell types to developmental, circadian and environmental cues. The Plant Journal 109: 764–778.

Taylor SC, Nadeau K, Abbasi M, Lachance C, Nguyen M, Fenrich J. 2019. The Ultimate qPCR Experiment: Producing Publication Quality, Reproducible Data the First Time. Trends in Biotechnology 37: 761–774.

Triviño M, Martín-Trillo M, Ballesteros I, Delgado D, de Marcos A, Desvoyes B, Gutiérrez C, Mena M, Fenoll C. 2013. Timely expression of the Arabidopsis stoma-fate master regulator MUTE is required for specification of other epidermal cell types. The Plant Journal 75: 808–822.

Vu LD, Xu X, Gevaert K, De Smet I. 2019. Developmental Plasticity at High Temperature. Plant Physiology 181: 399–411.

Wang H-Z, Yang K-Z, Zou J-J, Zhu L-L, Xie ZD, Morita MT, Tasaka M, Friml J, Grotewold E, Beeckman T, et al. 2015. Transcriptional regulation of PIN genes by FOUR LIPS and MYB88 during Arabidopsis root gravitropism. Nature Communications 6: 8822.

Wingett SW, Andrews S. 2018. FastQ Screen: A tool for multi-genome mapping and quality control. F1000Research 7: 1338.

Woodward FI. 1987. Stomatal numbers are sensitive to increases in CO2 from pre-industrial levels. Nature 327: 617–618.

Woodward FI, Lake JA, Quick WP. 2002. Stomatal development and CO _2_ : ecological consequences. New Phytologist 153: 477–484.

Xie Z, Lee E, Lucas JR, Morohashi K, Li D, Murray JAH, Sack FD, Grotewold E. 2010. Regulation of Cell Proliferation in the Stomatal Lineage by the Arabidopsis MYB FOUR LIPS via Direct Targeting of Core Cell Cycle Genes. The Plant Cell 22: 2306–2321.

Xue X, Bian C, Guo X, Di R, Dong J. 2020. The MAPK substrate MASS proteins regulate stomatal development in Arabidopsis. PLOS Genetics 16: e1008706.

Zhu T, Fonseca De Lima CF, De Smet I. 2021. The heat is on: how crop growth, development, and yield respond to high temperature (S Penfield, Ed.). Journal of Experimental Botany: erab308.

Ziegler H. 1987. The evolution of stomata. In: Stomatal function. Stanford University Press, 29–57.

Zuch DT, Herrmann A, Kim E-D, Torii KU. 2023. Cell Cycle Dynamics during Stomatal Development: Window of MUTE Action and Ramification of Its Loss-of-Function on an Uncommitted Precursor. Plant and Cell Physiology 64: 325–335.

Zuo J, Niu Q-W, Chua N-H. 2000. An estrogen receptor-based transactivator XVE mediates highly inducible gene expression in transgenic plants. The Plant Journal 24: 265–273.

